# Spontaneous calcium transients in hair cell stereocilia

**DOI:** 10.1101/2024.08.12.607658

**Authors:** Saman Hussain, Miloslav Sedlacek, Runjia Cui, Wendy Zhang-Hooks, Dwight Bergles, Jung Bum-Shin, Katie S. Kindt, Bechara Kachar

## Abstract

The hair bundle of auditory and vestibular hair cells converts mechanical stimuli into electrical signals through mechanoelectrical transduction (MET). The MET apparatus is built around a tip link that connects neighboring stereocilia that are aligned in the direction of mechanosensitivity of the hair bundle. Upon stimulation, the MET channel complex responds to changes in tip-link tension and allows a cation influx into the cell. Ca^2+^ influx in stereocilia has been used as a signature of MET activity. Using genetically encoded Ca^2+^ sensors (GCaMP3, GCaMP6s) and high-performance fluorescence confocal microscopy, we detect spontaneous Ca^2+^ transients in individual stereocilia in developing and fully formed hair bundles. We demonstrate that this activity is abolished by MET channel blockers and thus likely originates from putative MET channels. We observe Ca^2+^ transients in the stereocilia of mice in tissue explants as well as *in vivo* in zebrafish hair cells, indicating this activity is functionally conserved. Within stereocilia, the origin of Ca^2+^ transients is not limited to the canonical MET site at the stereocilia tip but is also present along the stereocilia length. Remarkably, we also observe these Ca^2+^ transients in the microvilli-like structures on the hair cell surface in the early stages of bundle development, prior to the onset of MET. Ca^2+^ transients are also present in the tallest rows of stereocilia in auditory hair cells, structures not traditionally thought to contain MET channels. We hypothesize that this newly described activity may reflect stochastic and spontaneous MET channel opening. Localization of these transients to other regions of the stereocilia indicates the presence of a pool of channels or channel precursors. Our work provides insights into MET channel assembly, maturation, function, and turnover.

## Introduction

In auditory and vestibular hair cells, mechanoelectrical transduction (MET) takes place in the hair bundle, a specialized organelle located on the apical surface of hair cells (Schwander et al., 2010). Each hair bundle consists of a collection of actin-filled membrane protrusions called stereocilia organized in a lengthwise order to form a staircase pattern. Shorter stereocilia are connected to their taller neighbor via a tip link composed of cadherin 23 and protocadherin 15 (Kazmierczak et al., 2007). The MET apparatus is built around the tip link with the upper end connected to a tension-generating motor (Grati and Kachar, 2011) and the lower end attached to the MET channel complex (Beurg et al., 2009). Mechanical stimuli can pivot the hair bundle in the direction of the tallest stereocilia, which increases tip link tension and results in the opening of the MET channel. This is followed by an influx of cations, mainly K^+^ and Ca^2+^ (Peng et al., 2011) which depolarizes the hair cell (Figure 1A).

**Figure 1:**
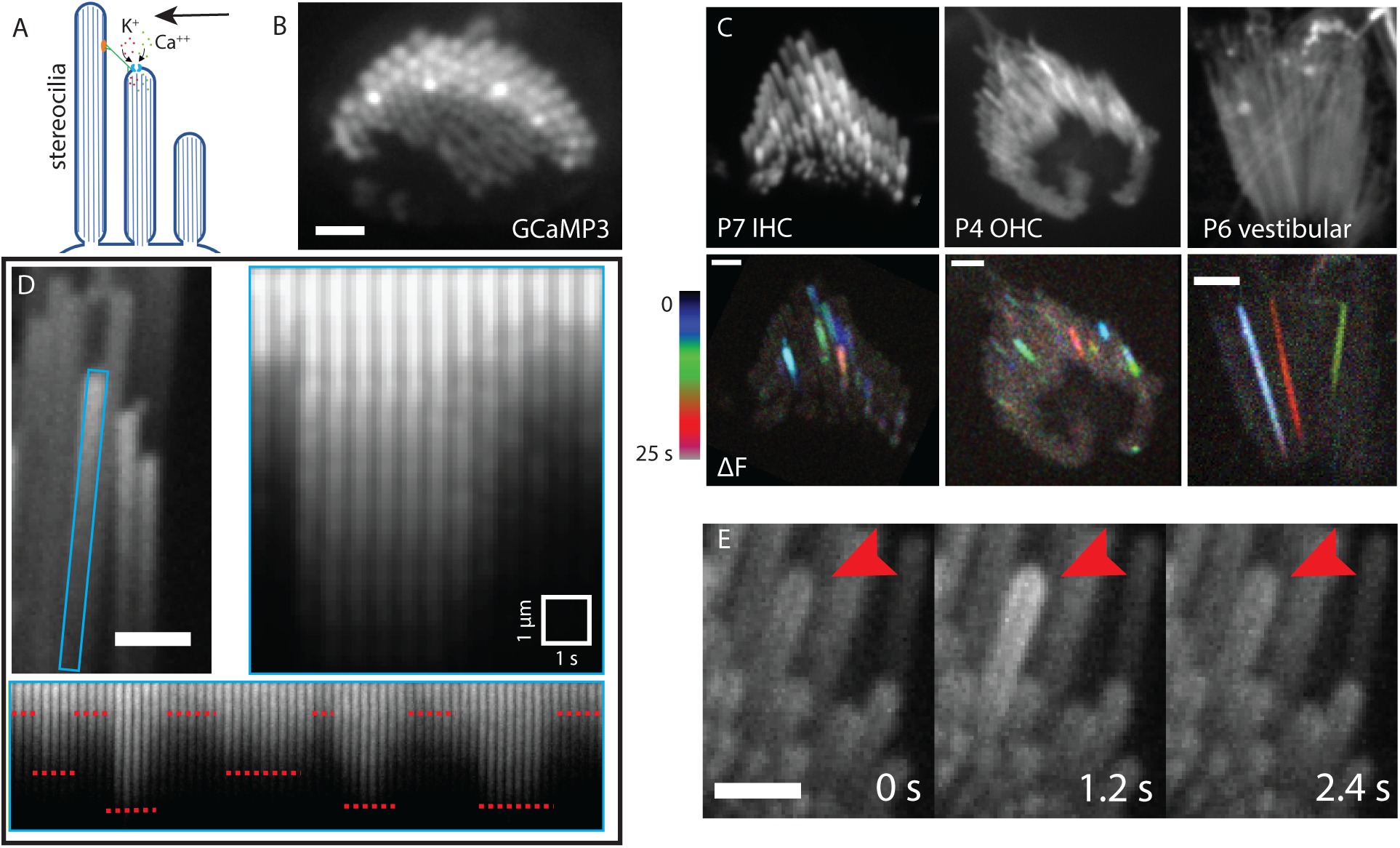
Characteristics and prevalence of spontaneous calcium transients. **A)** A diagrammatic representation of stimulated MET channel activity. When the hair bundle deflects (black arrow) there is an increased tension in the tip link (green). This increases the open probability of the MET channel and results in an influx of cations. **B)** A top-down view of a hair bundle with calcium transients in individual stereocilia, indicated by an increased GCaMP3 signal. Scale bar is 2 *μ*m. **C)** Temporal color-coded projections of spontaneous calcium transients are shown in a P7 inner hair cell (left), P4 outer hair cell (middle) and P6 vestibular hair cell (right). The transients do not appear to be spatially or temporally correlated. Scale bar is 2 *μ*m. **D)** The stereocilium marked by a blue rectangle is plotted over time, with a time interval of 500 ms. An individual calcium transient in this stereocilium is shown on the right panel. GCaMP3 signal increases sharply within a span of 500 ms and decays gradually over 3 s. The same stereocilium tip displays variable but step-like changes in intensity over a longer time period. The steps are marked by red dotted lines. Scale bar is 1 *μ*m. **E)** Montage of a calcium transient over time in a P6 inner hair cell with GCaMP3. Red arrowhead points to the stereocilium displaying the activity. Each frame is 1.2 s apart. Scale bar is 2 *μ*m.

Understanding the MET channel gating mechanisms comes primarily from electrophysiological experiments. It is known that in an unstimulated state, the tip link experiences a resting tension, which modulates the resting open probability of the MET channel and results in a small overall resting hair cell current (Corey and Hudspeth, 1979). However, the precise nature or physiological significance of this baseline activity is not known. There is evidence that Ca^2+^ influx through the MET channel may affect actin remodeling and help regulate the length and morphology of hair bundles (Velez-Ortega et al., 2017). Furthermore, resting MET currents at the pre-hearing stage are likely required for proper hair bundle development and functional maturation (Corns et al 2018).

Ca^2+^ imaging has been used previously to study MET channel activity and localization during hair bundle stimulation (Beurg et al., 2009). However, spontaneous MET channel activity in unstimulated bundles has not been reported, likely due to the challenges in detection related to the limited spatiotemporal resolution of Ca^2+^ imaging approaches, which make it difficult to resolve individual stereocilia (∼ 200 nm diameter). In addition, spontaneous MET channel activity is likely to occur at a rather low frequency and with low signal intensity as it depends on a single or a discrete number of channels opening at a time (Beurg et al., 2018). Here, by performing Ca^2+^ imaging using genetically encoded membrane bound Ca^2+^ sensors GCaMP3 and GCaMP6s, and high-performance confocal microscopy, we report localized spontaneous Ca^2+^ transients in individual stereocilia mediated by MET channels at rest. This spontaneous activity was reliably detected both in developing and mature hair bundles of mice. Spontaneous Ca^2+^ transients were observed in individual stereocilia in hair bundles from the earliest post-natal stages (P0) and continued to occur in later developmental stages (after the onset of hearing). In addition, spontaneous Ca^2+^ transients were also observed in the individual stereocilia of zebrafish hair cells, indicating that this activity is conserved and present *in vivo*. By using pharmacological tools, we confirm that this activity is associated with MET channels, and we demonstrate that membrane tension can play a role in triggering this activity. Furthermore, we show that the origin of these Ca^2+^ transients is not limited to the canonical MET site at the stereocilia tip but is also observed along the stereocilia length. We report the presence of these transients in previously unknown locations including the microvilli on the apical surface of developing hair cells, and the tallest rows of stereocilia. We discuss the likely significance of this spontaneous activity to the process of MET channel assembly, maturation, function, and turnover.

## Results

### Individual stereocilia exhibit spontaneous Ca^2+^ transients

To determine if there was spontaneous Ca^2+^ activity in unstimulated hair bundles, we used a transgenic mouse line that expresses a membrane-bound calcium indicator GCaMP3 selectively in hair cells (Agarwal et al., 2017). The baseline GCaMP3 fluorescence in this line allowed us to visualize stereocilia at resting intracellular Ca^2+^ levels. By imaging GCaMP3 signals in stereocilia at an acquisition rate of 1 to 5 frames per second, we observed Ca^2+^ transients in individual stereocilia of early postnatal mice (P4 – P9) in the absence of any external mechanical stimulation (Figure 1, Supplemental Movie 1). We observed Ca^2+^ transients in the stereocilia of both inner and outer auditory hair cells of the organ of Corti (Figure 1C, Supplemental Movie 2). In addition, when we expanded our GCaMP3 imaging to the vestibular system, we also observed spontaneous Ca^2+^ transients in the stereocilia of utricular and saccular hair cells (Figure 1C, Supplemental Movie 2). We found that Ca^2+^ transients in each individual stereocilium occurred independently of Ca^2+^ transients in neighboring stereocilia. To visually demonstrate that neighboring events were temporally uncorrelated, GCaMP3 signals were displayed as temporal color-coded projections (Figure 1C). We also found that Ca^2+^ transients were not spatially clustered within the hair bundle, which might be expected if the Ca^2+^ transients were occurring in response to deflection of stereocilia by local or global fluid perturbations in the sample. Next, we expanded our imaging to older ages to understand if these Ca^2+^ transients persist even after the onset of hearing. For this analysis, we imaged hair cells in acute tissue whole mounts of P17 organs of Corti. Using this approach, we were able to detect spontaneous Ca^2+^ transients, indicating that this activity persists even after the MET complex is fully mature (Supplementary Figure 1).

## The amplitude and duration of spontaneous Ca^2+^ transients are variable

After identifying spontaneous Ca^2+^ transients in the stereocilia of mouse auditory and vestibular hair cells, we examined the amplitude and duration of these transients more closely. For amplitude measurements, we analyzed the distance each Ca^2+^ transient propagated from its point of origin. For this analysis, we created an ROI of the individual stereocilium (blue ROI in Figure 1D). We then plotted a line along the center of the individual stereocilium and generated a kymograph of GCaMP3 signals along this line over time. We found that the propagation distance varied for each Ca^2+^ transient, even within the same stereocilium (Figure 1D, bottom panel). Interestingly, our results also revealed that distinct Ca^2+^ transient amplitudes were repeated over time (Figure 1D, bottom panel). This variable, yet repeated, amplitude can be explained by two mechanisms. It is possible that 1) multiple MET channels exist within a complex and different number of channels are open in each state, a possibility that has been suggested previously in electrophysiological experiments (Beurg et al., 2018), or 2) the MET channel is a single channel with varying conductance levels (Beurg et al., 2015).

We also extended our analyses to quantify the temporal components of the Ca^2+^ transients. Within stereocilia, we observed variable temporal profiles with regard to both rise time and decay. For example, in some instances the total rise time occurred within a single acquisition frame (Supplementary Figure 1B), while in other instances it took 3 - 5 seconds for the transients to peak (Figure 1D). It is important to note that MET speed far exceeds the GCaMP3 kinetics as well as the speed of our recordings, so we are limited in temporal resolution when interpreting the kinetics of channel gating. Similar variability was observed in the decay of the Ca^2+^ transients. In the majority of Ca^2+^ transients, the decrease in GCaMP3 fluorescence occurred within 1-5 seconds (Figure 1D). Occasionally, we observed a slow gradual decrease in GCaMP3 fluorescence lasting several minutes (Supplementary Figure 1B). The variability in decay rate is most likely a function many factors including internal Ca^2+^ buffers and Ca^2+^ extrusion from the stereocilia via PMCA2 (Dumont et al., 2001, Lumpkin and Hudspeth, 1998, Lumpkin and Hudspeth, 1995, Yamoah et al., 1998). In addition, the kinetic profile of the Ca^2+^ indicator likely impacts the temporal profile of the Ca^2+^ transients as well (Beurg et al., 2009). Overall, our imaging of Ca^2+^ transients indicates that transients are highly variable in duration and propagation, even within an individual stereocilium.

### Ca^2+^ transients are sensitive to MET channel blockers

Given the location of Ca^2+^ transients in stereocilia, MET channel opening is the likely source of this spontaneous activity. To confirm that MET channels are required for spontaneous Ca^2+^ transients, we used the well-established MET channel blocker dihydrostreptomycin (DHS, 200 µM and 1 mM) (Kenyon et al., 2021, Kimitsuki and Ohmori, 1993). For this pharmacological analysis, we recorded GCaMP3 signals in auditory and vestibular tissue explants treated with DHS and compared these measurements to those captured in untreated control explants. Compared to controls, we found that 1 mM DHS-treated explants displayed a dramatic decrease in the number of Ca^2+^ transients in stereocilia, with most time-lapse recordings showing no transients at all (Figure 2A). We examined individual hair cells and counted the number of active stereocilia per hair cell. We saw a significant, dose-dependent reduction in the number of stereocilia with Ca^2+^ transients in samples treated with both 200 μM (mean = 17.1, n = 9 hair cells) and 1 mM (mean = 8.33, n = 10 hair cells) DHS compared to the untreated control (mean = 0.6, n = 10 hair cells) (Figure 2B).

**Figure 2:**
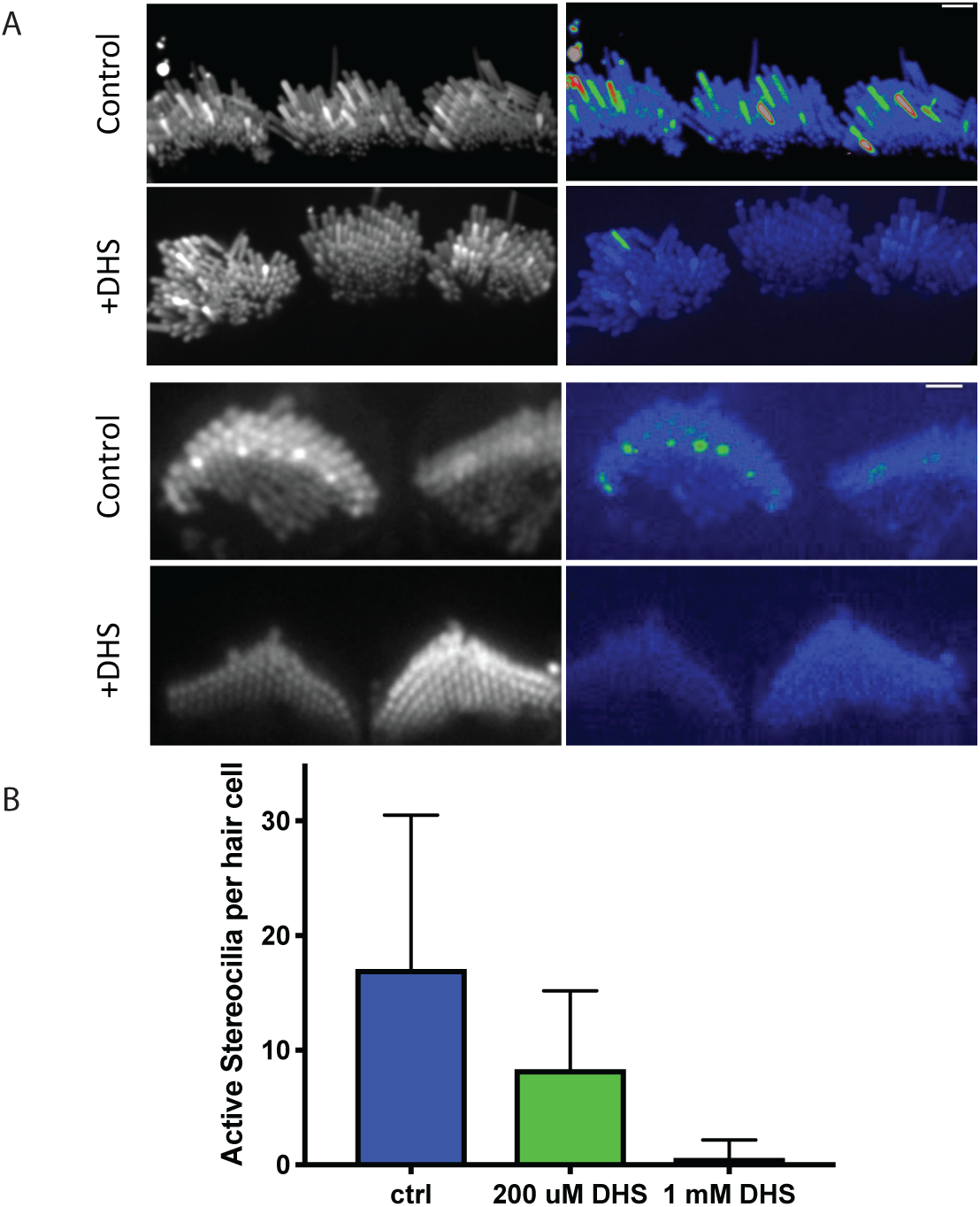
Spontaneous calcium transients are greatly reduced upon treatment with the mechanotransduction (MET) channel blocker dihydrostreptomycin (DHS). **A)** Shown here are standard deviation projections of cultured P5 organs of Corti in control and 1 mM DHS-treated cells. The untreated stereocilia exhibit more spontaneous activity and hence show up brighter in the standard deviation images (right column). Scale bars are 2 *μ*m. **B)** The number of active stereocilia per hair cell were counted by visual inspection and the numbers recorded. Cultured P4 organs of Corti were used in the quantification, and n = 10, 9 and 10 hair cells for the control, 200 *μ*M, 1 mM conditions respectively. Error bars are SD.

In addition, we confirmed this result by using another MET channel blocker, amiloride and saw that Ca^2+^ activity in stereocilia was drastically reduced by 50 μM amiloride treatment (Supplemental Movie 3). Together, our pharmacological data strongly indicate that the spontaneous Ca^2+^ transients most likely originate from MET channel activation.

### Ca^2+^ transients originate primarily at or near stereocilia tips

After confirming that MET channels were required for spontaneous Ca^2+^ transients, we examined where on stereocilia Ca^2+^ events originate. As MET channels are localized primarily at the tip of stereocilia, we hypothesized that this is where spontaneous events may originate. The spatial and temporal resolution of our imaging approach was sufficient to follow the onset and progression of Ca^2+^ transients along individual vestibular hair cell stereocilium. For this analysis we plotted the intensity along a line within the stereocilium. When we examined the first frame a transient was detected, we found that most Ca^2+^ transients originated at or near the tips of the stereocilia, the primary location of MET channels (Example, Figure 3A, B). Interestingly, we also observed that the origin of the Ca^2+^ transient could vary, and transients initiated in various locations along the stereocilium. We even observed variability in onset location within the same stereocilium (Figure 3C).

**Figure 3:**
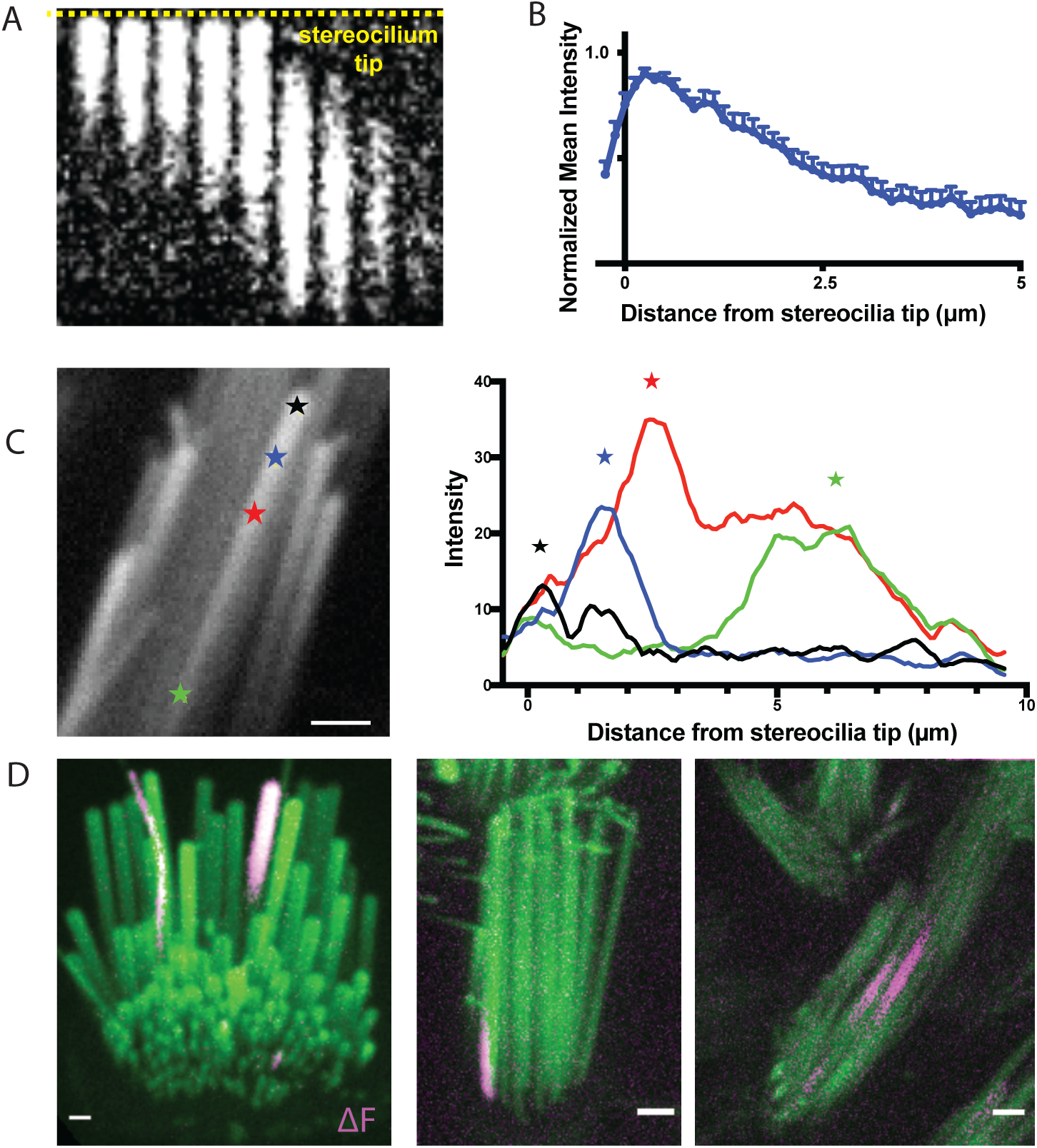
Transients predominantly originate near the stereocilia tip but are occasionally observed in the middle and basal regions. **A)** A kymograph of a transient is shown. The yellow dashed line marks the stereocilium tip. Transients predominantly originate near the stereocilium tip. Frames are 400 ms apart. **B)** Line intensity profiles along the length of 15 stereocilia were measured at the first frame where a transient was seen. The values were normalized to the maximum intensity of each trace. The mean and SEM are plotted here. The resulting graph shows that the origin of the transients is predominantly at the stereocilia tip. **C)** Within the same stereocilium, transients can originate from di]erent locations as shown in this example. The line intensity profiles along the length of this stereocilium are plotted at the first frame a calcium transient is seen for 4 transients. The transients originate from di]erent points along the length of the stereocilium, marked by stars. **D)** Transients are occasionally seen to originate from unexpected locations such as (i) in the tallest row in the organ of Corti where the MET channel is not believed to be present (right, shown in a P5 inner hair cell), (ii) at the base of stereocilia (middle, shown in a P9 vestibular hair cell), and (iii) in the middle of stereocilia (right, shown in a P4 vestibular hair cell). Scale bars are 1 *μ*m.

Occasionally, we observed transients that originated at a significant distance from the stereocilia tip, such as the middle portion and the base of vestibular stereocilia (Figure 3C (green star), 3D (middle and right panel), Supplemental Movie 4 (second and third movie)). These rare base transients do not appear to be temporally linked to cell body depolarization or Ca^2+^ transients in neighboring stereocilia. It is possible that the spontaneous events initiating closer to the base of stereocilia originate from the MET channel while it is trafficked to the stereocilia tips during bundle development or during channel complex turnover (Kachar et al., 2000, Indzhykulian et al., 2013). In addition, this activity may arise from a pool of MET channels or channel complex precursors, which have been reported to have a stochastic distribution along stereocilia during development (Kurima et al., 2015). Another possibility is that these stereocilia base transients originate from non-canonical MET channels such as Piezo2 which have been reported to localize near the base of stereocilia (Wu et al., 2017, Beurg et al., 2016).

It is currently accepted that the MET channel complex is present at the lower end of the tip links (Beurg et al., 2009). In mature auditory hair cells this arrangement means that the second and third row of stereocilia are mechanosensitive, while the tallest row is not (Beurg et al., 2009, Schwander et al., 2010). However, in auditory hair cells, we also observed spontaneous Ca^2+^ transients occurring in the tallest row in mice ranging from P2 to as old as P17 (Figure 3D, Supplemental Movie 4 (first movie)). Importantly, events in the tallest stereocilia occurred even when the adjacent, shorter stereocilia coupled by a tip link had no Ca^2+^ event. This finding is not in line with the idea that MET channels are absent from the upper end of the tip link. It is possible that a pool of MET channels or channel complex can persist on the tips of the tallest row of stereocilia (Kurima et al., 2015). These channels may not have a canonical role in hair cell mechanotransduction.

### Ca^2+^ transients in developing stereocilia precede the onset of mechanotransduction

In auditory hair cells, we have found that spontaneous Ca^2+^ transients in stereocilia occur at very early ages (P0-P3) (Figure 4A). These ages are prior to the onset of canonical mechanosensitivity in mouse auditory hair cells (Waguespack et al., 2007). At these early ages, the apical surface of hair cells is covered with microvilli-like structures (Figure 4B). Some of these microvilli elongate and form mature stereocilia, while others, called supernumerary microvilli, shrink back into the apical membrane (Krey et al., 2023, Tilney et al., 1992, Schwander et al., 2010). In auditory hair cells from P0-P3 mice, the majority of Ca^2+^ transients we observed were in taller microvilli that are maturing into stereocilia. However, we also saw spontaneous Ca^2+^ transients in the microvilli or stereocilia precursors that are not yet part of the emerging staircase-shaped stereocilia bundle (Figure 4B, red arrowheads, Supplemental Movie 5). We observed this activity in 3 out of 11 hair cells with microvilli. In those hair cells that displayed spontaneous calcium activity in microvilli, this activity occurred at an average frequency of 1.6 active microvilli per hair cell. Interestingly we observed that the time course of Ca^2+^ transients in the precursors and in stereocilia are similar (Figure 4C). These microvilli are not sites where canonical MET current is presumed to occur. But this observation does match previous studies detecting MET channel complex components TMC1 and TMC2 in these microvilli (Kurima et al., 2015).

**Figure 4:**
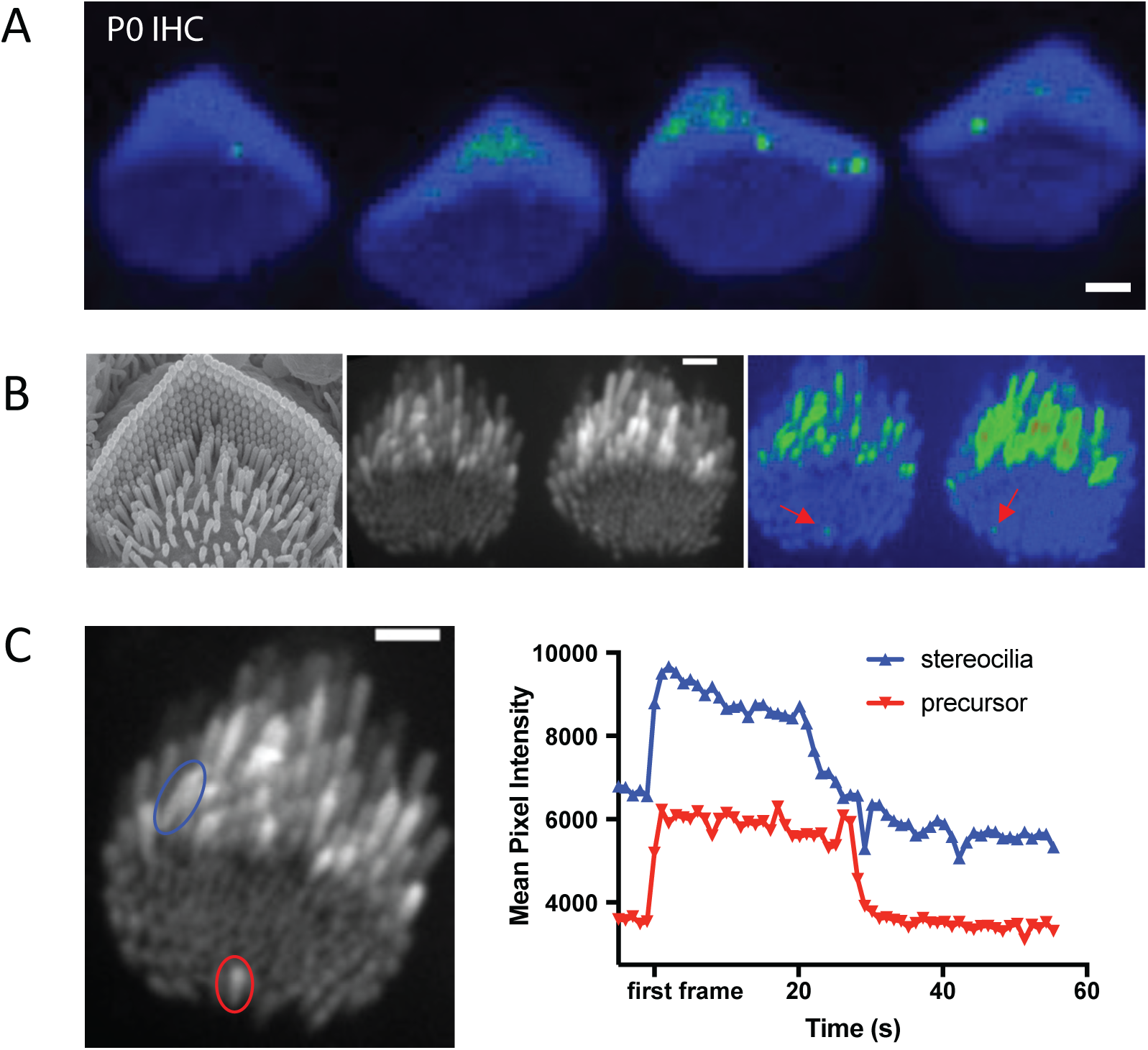
Calcium transients in stereocilia precede the maturation of the MET complex. **A)** A standard deviation projection of a time-lapse that shows spontaneous calcium transients occurring in stereocilia inner hair cells at P0, prior to the maturation of the MET complex. **B)** Immature microvilli cover the apical surface of early post-natal hair cells as shown in the SEM image *(left).* Only a few of these microvilli precursors elongate to become stereocilia while the rest disassemble and are integrated into the cell membrane. These microvilli also exhibit spontaneous calcium transients as shown in the standard deviation projection (*right*) of P4 inner hair cells, although their occurrence is much rarer than those in stereocilia. **C)** Transients in microvilli precursors occur on the same timescale as in stereocilia. Shown are traces for a stereocilium (blue) and a precursor (red). The timeframes for the traces have been shifted such that t = 0 when the transient is first seen.

### Stereocilia Ca^2+^ transients can occur in the presence or absence of somatic Ca^2+^ signaling

Spontaneous Ca^2+^ activity in the hair cell soma has been extensively studied in previous work. In the developing cochlea, purinergic signaling among supporting cells can induce somatic Ca^2+^ activity in collections of hair cells. This coordinated activity is essential for the proper formation of the auditory neural circuit (Kersbergen et al., 2022, Kersbergen and Bergles, 2024, Sun et al., 2018). However, hair cells also exhibit spontaneous, somatic Ca^2+^ activity that does not require purinergic signaling and is not correlated with the activity of neighboring hair cells ((Wang and Bergles, 2015); Supplemental movie 6). To determine if somatic Ca^2+^ activity is correlated to the activity in individual stereocilia, we acquired 3D stacks of vestibular hair cells (ampullar). We mounted and imaged the cells at an angle where both the cell soma and the hair bundle were present in the field of view. In our 3D stacks, we observed Ca^2+^ transients occurring in stereocilia of hair cells that both exhibited or did not exhibit Ca^2+^ transients in their soma (Figure 5A, Supplemental Movie 7). This suggests that spontaneous activity in the soma is independent of the spontaneous Ca^2+^ transients in the stereocilia. To further confirm that these Ca^2+^ transients in the soma and stereocilia were independent, we used cytoplasmic GCaMP6F to measure Ca^2+^ transients in the hair cell soma. Compared to controls, we found that spontaneous Ca^2+^ transients in the soma continued to occur in these cells after 1 mM DHS exposure (Figure 5B). In our previous DHS manipulation, we observed that the majority of stereocilia Ca^2+^ events abolished after DHS exposure (Figure 3A-B). Together these DHS experiments strongly indicates that spontaneous Ca^2+^ transients in the soma and stereocilia occur independently, and that these transient MET openings do not provide enough depolarization to trigger calcium action potentials.

**Figure 5:**
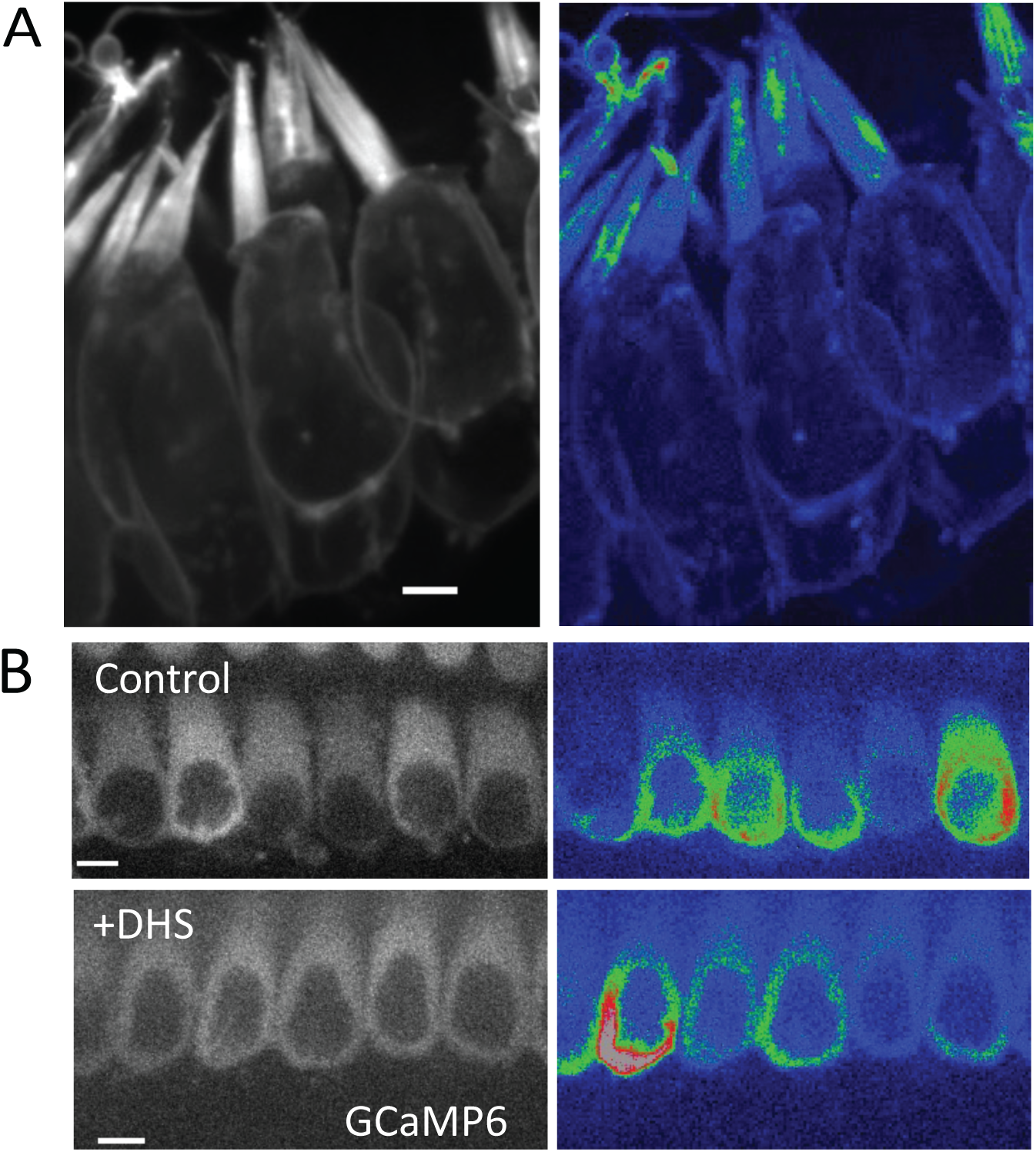
Calcium transients in stereocilia appear to be independent of whole cell spontaneous activity. **A)** An example of P0 ampullar hair cells containing stereocilia with calcium transients but no spontaneous activity in the cell body. An example of cells exhibiting cell body activity and calcium transients in stereocilia is shown in Supplemental Movie 7. Scale bar is 5 *μ*m**. B)** DHS treatment (1 mM) does not a]ect the spontaneous activity in the hair cell body, which is controlled by purinergic signaling and does not require a functional MET channel. Cultured P2 organs of Corti are shown here with cytoplasmic expression of GCaMP6F. Scale bar is 5 *μ*m.

### Membrane tension and tethering can trigger Ca^2+^ transients

To image spontaneous Ca^2+^ signals in stereocilia, we were careful to ensure that the hair bundles were not under tension. We observed that when either auditory or vestibular sensory tissues were gently pressed onto the microscope coverslip, the hair bundles made contact with the glass surface and became over-deflected. Interestingly this persistent deflection was also accompanied by the formation of membrane tethers, very thin protrusions from the tips of the stereocilia. The tethers formed between adjacent stereocilia as well as between a stereocilium and the glass coverslip (Figure 6A *arrows*, Supplemental Movie 8). We observed Ca^2+^ transients originating from the tips of these membrane tethers (Figure 6B, Supplemental Movie 8).

**Figure 6:**
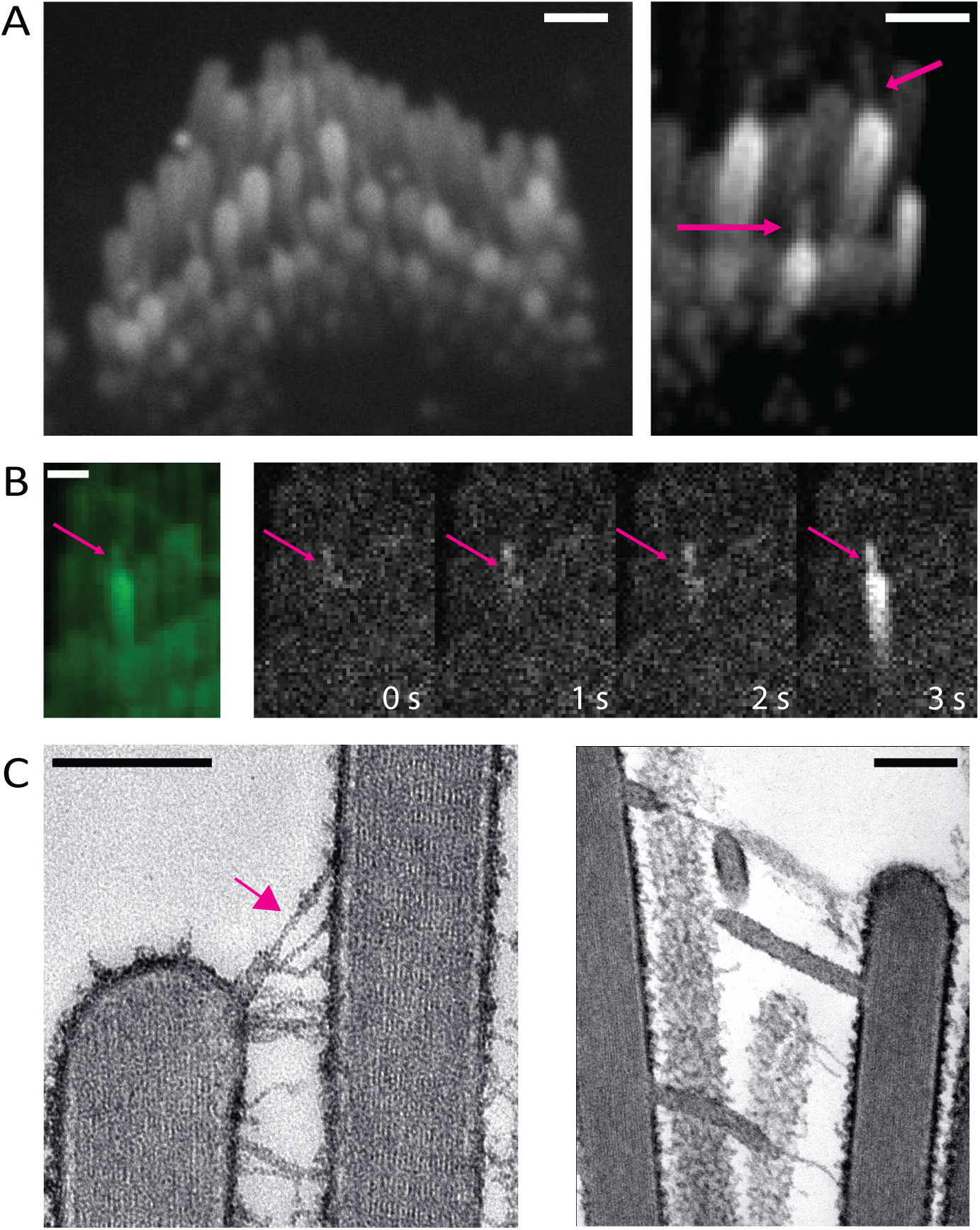
Tension in membrane tethers could play a role in generating calcium transients in stereocilia. **A)** Over-deflected stereocilia experience a shear force against the coverslip. This results in the formation of membrane tethers which can be visualized with membrane-localized GCaMP3. The tethers can occur between two stereocilia or between the glass surface and a stereocilium (*magenta* arrows). A large proportion of the tethered stereocilia appear bright, indicating an increased calcium influx. Scale bars are1 *μ*m. **B)** Membrane tethers play a role in calcium influx, and tethers can act as sites for the origination of spontaneous calcium transients. The intensity di]erence time series shown here was taken every 1s. It can be observed in this example that the membrane tether (magenta arrow) shows an increased calcium level first, followed by a calcium transient in the stereocilium. Scale bar is 1 *μ*m. **C)** Over-deflected hair bundles in electron microscopy preparations display membrane tethers. The tip link remains intact at the tether end (magenta arrow, left). Scale bar is 200 nm.

To investigate the ultrastructure of these membrane tethers, we examined directly frozen vestibular hair bundles by thin section electron microscopy. Freshly dissected vestibular sensory epithelia were directly frozen by the rapid contact-freezing method and processed for freeze-substitution and low temperature embedded to avoid lipid solvents and better preserve membranes (Bullen et al., 2014). Conveniently, the freezing process involves rapid impact of the surface of the sensory epithelium on the surface of a copper block cooled to liquid nitrogen temperature. This rapid freezing results in hair bundles that are frozen at various degrees of deflection. In the bundles that have been deflected beyond the elastic range of the inter-stereocilia links, we observed the formation of membrane tethers at the ends of tip links and lateral links (Figure 6C). Remarkably, the tethers were uniform in diameter (∼30 nm), independent of their length. We also found that the tip link insertion to the membrane is preserved adjacent to the membrane tether (arrow, Figure 6C). Based on tip link preservation, we hypothesize that some of the transmembrane or cytoplasmic components of the MET channel complex remain attached to the membrane tether, in particular the membrane components of the MET channel complex. The TEM images support the possibility that the Ca^2+^ transients we see at the tips of the membrane tethers result from gating of tip link-associated MET channels that get pulled away during the tethering process and are maintained under tension by the elastic properties of the membrane tether (Nussenzveig, 2018, Pontes et al., 2013). This also indicates that membrane tension or pre-stress could be playing a role in gating the channel in the absence of a dynamic deflection of the bundle and that is why we see frequent Ca^2+^ transients at the tips of tethers.

### Stereocilia Ca^2+^ transients can be observed *in vivo* in zebrafish hair cells

In addition to cultured mouse organs of Corti and vestibular tissue, we wanted to investigate if spontaneous Ca^2+^ transients in individual stereocilia also occur in intact tissue *in vivo*. This was to ensure that our observations were not artifacts caused by the *in vitro* culturing of the mouse explants. To measure Ca^2+^ transients *in vivo*, we examined zebrafish lateral line and inner ear hair cells. We used transgenic zebrafish lines that expresses a membrane-localized, hair-cell specific GCaMP6 (m6b::GCaMP6s-caax). Compared to mice, the hair bundles in zebrafish are smaller and imaging stereocilia is more challenging. Therefore, to resolve Ca^2+^ transients in individual stereocilia in zebrafish, we used Zeiss Airyscan confocal microscopy. This approach enabled us to image Ca^2+^ transients at a rate and a resolution sufficient to resolve individual stereocilia and image the transients. We first examined hair cells within the lateral line. The lateral line is made up of clusters of hair cells called neuromasts (Example neuromast, Figure 7A). We examined GCaMP6s signals at 2 days post fertilization (dpf) when the hair bundles are developing, and at 5 dpf, when most hair bundles are mature (Kindt et al., 2012, Trapani and Nicolson, 2011, McHenry et al., 2009). We observed spontaneous Ca^2+^ transients in the stereocilia at both 2 and 5 dpf (Figure 7 A-B, Supplemental Movie 9). We extended our analysis to the cristae of the zebrafish inner ear and observed similar spontaneous Ca^2+^ transients in the stereocilia (Figure 7B, Supplemental Movie 9). This work in zebrafish indicates that spontaneous Ca^2+^ transients in stereocilia are conserved and are present *in vivo*.

**Figure 7:**
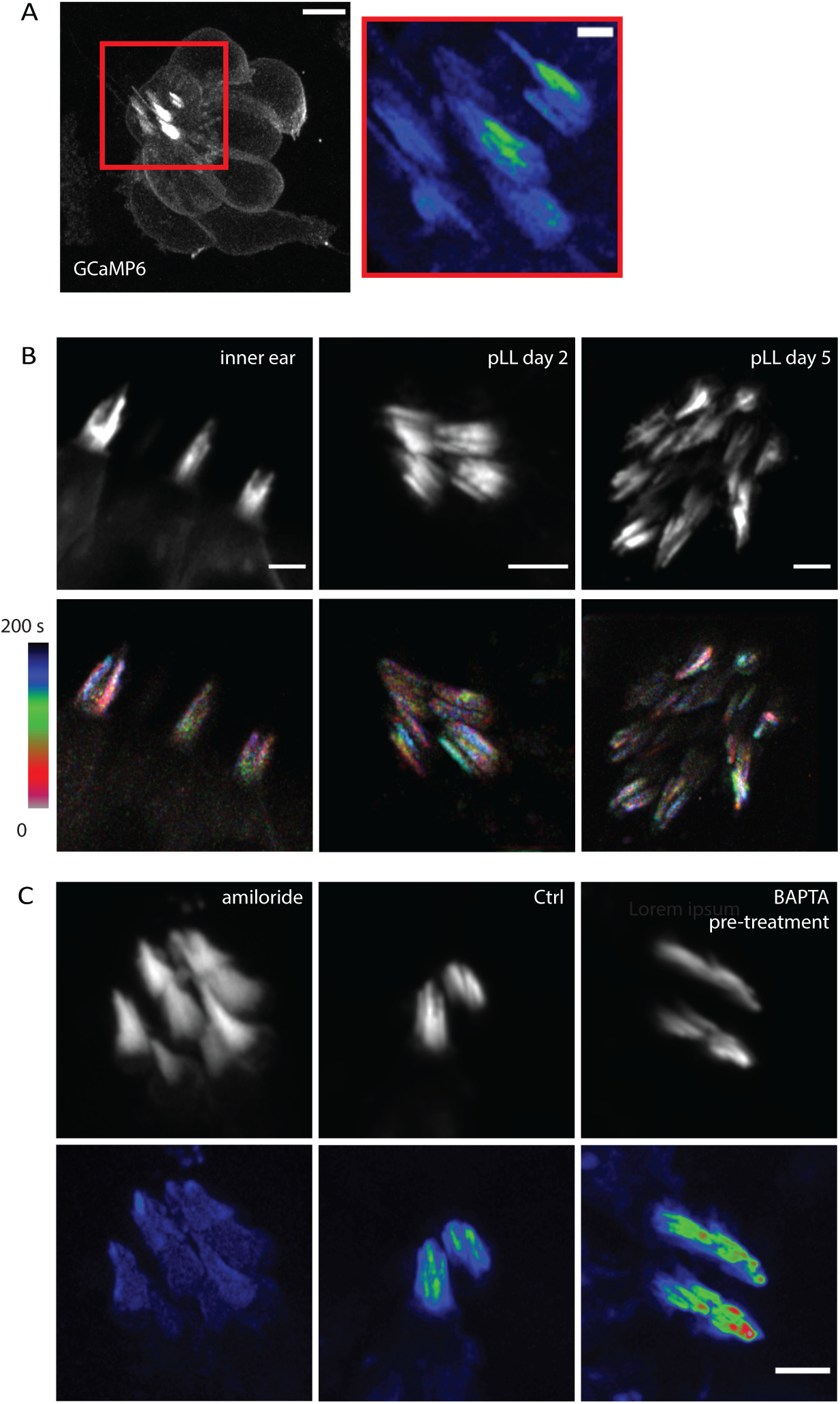
Spontaneous calcium transients are present *in vivo* in zebrafish hair cells. **A)** A 2 dpf zebrafish lateral line neuromast visualized with membrane-localized GCaMP6 is shown here. This neuromast was imaged every 16 s for 8 min to visualize calcium activity. The standard deviation projection of the region marked by the red rectangle is shown on the left. Calcium activity in stereocilia can be observed in the projection. Scale bar is 5 *μ*m (left image) and 1 *μ*m (right image). **B)** Calcium activity in stereocilia is present in the inner ear, in developing hair bundles at 2 dpf, and in mature hair bundles at 5 dpf. Shown in the top panel are hair bundles visualized by membrane-localized GCaMP6. The bottom panel shows temporally color-coded images of these hair bundles, where the bundles were images every 5 s. It can be seen from these images that the calcium activity is uncorrelated between neighboring stereocilia. Scale bars are 2 *μ*m. **C)** 2 dpf lateral line hair cells treated with amiloride (1 mM) and pre-treated with BAPTA (5 mM for 15 min) are shown. Amiloride treatment abolishes the calcium activity while BAPTA pre-treatment does not a]ect these transients. The hair bundles were imaged every 5 s for 8-10 min. Scale bar is 2 *μ*m.

Similar to our work in mice, we tested whether MET channels were required for this activity using the MET channel blocker amiloride, which has previously been used in zebrafish (Trapani and Nicolson, 2011). We found that treatment with 1 mM amiloride blocked the spontaneous Ca^2+^ transients completely (Figure 7C). Additionally, we tested whether intact tip links were needed for these Ca^2+^ transients by treating lateral line hair cells with 5 mM BAPTA for 15 min, followed by a wash before imaging. It has been previously shown that this treatment is sufficient to break all tip links (Kindt et al., 2012). Our results revealed that BAPTA pre-treatment to break tip links did not affect the spontaneous Ca^2+^ transients (Figure 7C). Together, our pharmacological findings in zebrafish confirm that MET or MET-like channels are the source of the spontaneous activity, but these channels do not require an intact tip link to exhibit this activity. This result is consistent with our observation in mice, where splayed hair bundles with broken tip links continue to display spontaneous activity.

## Discussion

In the auditory hair cells of mice, MET develops during the first postnatal week. During this period, there are modifications in the organization of the hair bundle, including elongation of the stereocilia, formation of the staircase shape, and reabsorption of supernumerary microvilli (Kurima et al., 2015, Schwander et al., 2010). During the first week, the magnitude of the MET current increases to reach mature levels well before the onset of hearing (Waguespack et al., 2007), which occurs after the second postnatal week. How the MET current is produced prior to the onset of hearing, and what mechanisms underlie the organization of the MET channel complexes at their functional site are not known. Single MET channel activity can be observed by patch-clamping following mechanical stimulation under some experimental recording conditions. To our knowledge, there has been no reports of direct visualization of spontaneous single channel Ca^2+^ activity in stereocilia.

In the present study, we report the direct observation of spontaneous Ca^2+^ transients in individual stereocilia. This spontaneous Ca^2+^ transients are inhibited by MET channel blockers (dihydrostreptomycin and amiloride) suggesting that they are generated by MET channels and/or MET channel precursors. The described Ca^2+^ transients most likely reflect the stochastic opening of MET channels that produce the small resting whole cell currents recorded in the absence of mechanical stimulation (Corey and Hudspeth, 1979). This stochastic opening may be an intrinsic property of the MET channel related to its high intrinsic open probability, evidenced by the fact that the activity continues even when tip links are disrupted by BAPTA pre-treatment.

Although Ca^2+^ transients occur in the absence of dynamic deflection of stereocilia, we also see in these preparations that membrane pre-stress or tension can increase the probability of their occurrence. A close view of some of the slightly splayed bundles show that Ca^2+^ transients originate more frequently at the sites where there is formation of membrane tethers. Tether formation requires a pulling force on the membrane. Tethers are also characterized by sustaining an elastic tension on the membrane (Nussenzveig, 2018, Pontes et al., 2013, Cugno et al., 2021, Li et al., 2002).

Ca^2+^ influx at the sites where membrane tension resulted in tether formation could in principle cause local ruptures in the membrane lipid bilayer creating transient pores as discussed in other systems (Eremchev et al., 2023). Additionally channels such as Piezo 2 under static membrane tension (Jia et al., 2016) may be contributing to the calcium activity. However, because Ca^2+^ transients were abolished by DHS and amiloride, even in stereocilia containing membrane tethers, we conclude that the source of these Ca^2+^ transients is primarily MET channels. In this case the greater mebrane tension produced by the tethers could increase the open probability of the MET channel and explain the higher frequency of spontaneous events at tether sites (Ricci et al., 2006, Ricci and Kachar, 2007, Jia et al., 2016).

By reconstitution of purified TMC1/2 in proteoliposomes, it has been shown that the pore-forming unit of the MET complex is TMC 1/2 (Jia et al., 2020). The channel activity in this reconstituted system shows single opening events, which differ from the multi-channel events observed *in vivo* (Beurg et al., 2018). Fluorescently tagged TMC 1/2 at the lower tip link density also photo bleaches in multiple steps (Kurima et al., 2015). Given these pieces of evidence, the MET complex most likely consists of multiple channels which come together to form a super-complex in the native state. Thus, reconstitution alone cannot provide us with information about the native state of the MET complex. In addition, although electrophysiological recordings have been made by stimulating single stereocilia, there is no information about the resting state activity of individual MET channels (Ricci et al., 2006, Ricci and Kachar, 2007, Peng et al., 2011).

We observe calcium transients in the tallest row stereocilia of outer and inner hair cells, where MET channels are not believed to be present. We attribute this observation to the presence of MET channel/precursor pools in the tallest row stereocilia that do not play a direct role in MET. Alternatively, these channels could be involved in an unrecognized form of mechanotransduction by sensing local plasma membrane tension by linkage or attachment (Verpy et al., 2011) to the overlaying tectorial membrane. Pushing or pulling forces at the point of contact or attachment of the stereocilia to the tectorial membrane during fluid movement could activate the MET channels in the tallest row. Since the tectorial membrane is removed in the experimental setup for *in vitro* studies, mechanosensitivity would not be present in the tallest row if it occurs by linkage to the tectorial membrane.

We also report the presence of Ca^2+^ signals in other unusual and previously unrecognized locations such as in stereocilia precursors and P0 tissue in which no MET currents have been detected electrophysiologically (Waguespack et al., 2007). Moreover, we observe Ca^2+^ transients originating under membrane tension in tethers, which may be an unrecognized form of mechanotransduction in hair cells when they are overstimulated.

Overstimulation of the hair bundle when experiencing loud noises causes breakage of the tip links to prevent structural damage to the stereocilia (Osborne et al., 1988). These tip links can be repaired in avian (Zhao et al., 1996) (Kachar et al., 2000) and mammalian hair cells within 24 hours (Jia et al., 2009). The robust nature of tip link repair and recovery indicates that there may be a pool of MET components in stereocilia to facilitate efficient reassembly (Kurima et al., 2015, Velez-Ortega and Frolenkov, 2019, Indzhykulian et al., 2013, Kazmierczak et al., 2007). Electrophysiological studies can measure channel conductance but provide no information about where the channels are located, since stereocilia cannot be patched. By observing Ca^2+^ activity in the unstimulated bundles, we discovered the presence of spontaneous Ca^2+^ transients in regions other than the tip of stereocilia, such as the middle and base regions. These Ca^2+^ transients may be originating from a pool of MET components that are not fully mechanically gated yet but are poised to take over the crucial role at the stereocilium tip in the event of tip link breakage.

Spontaneous Ca^2+^ transients in stereocilia are short lived and are not sufficient to elicit whole cell depolarization. These Ca^2+^ transients are also uncorrelated to spontaneous somatic Ca^2+^ transients induced by supporting cell K^+^ release (Sun et al., 2018, Tritsch et al., 2010, Wang and Bergles, 2015). There is, however, electrophysiological evidence that knocking out components of the MET complex, such as TMIE, alters the pattern of spontaneous currents generated in the spiral ganglion neurons (Sun et al., 2018). This would imply that the MET complex does have an effect on the spontaneous activity pattern, but that may be through the whole bundle MET activity. We did not examine subtle changes to the ionic activity pattern of hair cells and only tested whether the cell body Ca^2+^ spiking is present or absent in hair cells treated with DHS and amiloride. Thus, it is possible that there are changes in the spontaneous activity pattern of the hair cell body when the spontaneous Ca^2+^ activity of stereocilia is inhibited.

Overall, our results indicate that stereocilia exhibit spontaneous calcium activity in an unstimulated state, and this activity occurs even when the tip links are broken. This activity is observed at various ages and in locations other than the stereocilia tips, showing that MET channels exhibit spontaneous gating in both developing and mature stereocilia. Thus, our work provides unique insight into the development, maturation and functionality of the MET complexes.

## ACKNOWLEDGEMENTS

This work was supported by National Institute on Deafness and Other Communication Disorders (NIDCD) Intramural Research Program Grant Z01-DC000002 and 1ZIADC000085-01.

## MATERIALS AND METHODS

### Ethics Statement

All experiments were conducted according to the Guide for the Care and Use of Laboratory Animals by the National Institute of Health and approved by the Animal Care and Use Committees for the National Institute on Deafness and Other Communication Disorders (NIDCD ACUC, Protocol #1215 and Protocol #1362-13).

### Transgenic Mice

Transgenic mice with membrane localized GCaMP3 (R26-lsl-GCAMP3) or cytoplasmic GCaMP6f on the C57BL/6 background were previously described (Agarwal et al., 2017) . These mice were bred to homozygosity and crossed with heterozygous Gfi1-Cre which resulted in hair-cell specific GCaMP3 expression in Cre positive offspring. Transgenic animals were genotyped by standard PCR using Cre primers and GCaMP3 primers.

### Sample preparation (Mice)

Organs of Corti and vestibular tissue were dissected in DMEM/F12 medium (Thermo Fisher Scientific) from post-natal mice (P0 – P20) at room temperature. The tissue was mounted on collagen-coated, sterilized glass coverslips or glass-bottomed MatTek dishes and placed in culture medium (DMEM/F12 containing 10% fetal bovine serum (FBS) and 5 μg/ml ampicillin). The samples were allowed to grow 24-48 hours in an incubator at 37°C in 5% CO_2._ The culture medium was replaced after 24 hours. Acute preparations of older organs of Corti (P17-P20) were used without culturing. For imaging on the inverted microscope, coverslips with mounted samples were inverted on a glass bottom dish and fixed using resin.

### Pharmacology (Mice)

P5-P7 organ of Corti samples were dissected and cultured as described in the previous section. Prior to imaging, the samples were treated with 200 μM or 1 mM dihydrostreptomycin (DHS), or 50 μM amiloride diluted in DMEM/F12 (Thermo Fisher Scientific). The samples were imaged after15-20 min incubation.

### Membrane tether formation (Mice)

To induce tether formation, the hair bundles must be over deflected. This was achieved by culturing the sample on coverslips and inverting the coverslip into a glass bottom dish. The coverslip was gently pressed down on a few spots of Vaseline used as spacers and for adhesion. This gentle pressure resulted in the contact of hair bundles with the surface of the glass and the splaying of the bundle with formation of tethers between stereocilia as well as between stereocilia and the glass-bottom dish.

### Microscopy (Mice)

Timelapses were obtained using a spinning disk confocal system on Nikon Ti inverted and upright microscopes equipped with a Yokogawa CSU-21 spinning disk system and an Andor DU-897 camera. A 488 nm laser was used for fluorescence excitation. Timelapses were acquired using Nikon Elements software.

### Zebrafish Husbandry and Strains

Zebrafish (*Danio rerio*) larvae were grown in E3 embryo medium (5 mM NaCl, 0.17 mM KCl, 0.33 mM CaCl_2_, and 0.33 mM MgSO_4_, pH 7.2) at 30 °C using standard methods by the Animal Use Committee at the NIH under animal study protocol #1362-13. Larvae were examined at 2–5 days post fertilization (dpf). We used a previously described transgenic zebrafish strain *Tg(myo6b:GCaMP6s-CAAX)^idc1Tg^* (Zhang et al., 2018).

### Sample Preparation (Zebrafish)

Individual larvae were anesthetized with tricaine (0.03% ethyl 3-aminobenzoate methanesulfonate salt, Sigma). The larvae were then mounted on to glass-bottom dishes (35 mm Iwaki, 9.380190; 27 mm ThermoScientific, 1141351). To immobilize the larvae, they were mounted in 1% low melting point agarose (Promega, V2111) solution made in E3 medium with added tricaine (at 0.02–0.04% w/v). Once the agarose set, saturated Kimwipes (Kimtech, 34155) with E3 media containing tricaine (at 0.02–0.04% w/v) were placed around the edges of the dish in a circle to prevent the agarose from drying out and the dish was sealed with parafilm.

### Treatment with amiloride and BAPTA (Zebrafish)

For amiloride treatment, zebrafish larvae at 2 dpf were mounted in 1% LMP agarose solution made in E3 medium with added tricaine (at 0.02–0.04% w/v) and 1 mM amiloride (A7410-5G, Sigma).

For BAPTA pre-treatment, zebrafish larvae at 2 dpf were placed in 5 mM BAPTA (B22258-06, ThermoScientific) solution made in E3 medium for 15 minutes. The BAPTA was then washed out and the larvae were mounted in 1% LMP agarose solution made in E3 medium with added tricaine (at 0.02–0.04% w/v).

### Microscopy (Zebrafish)

Zebrafish larvae were imaged on the Zeiss LSM 980 system with Airyscan using a 63x oil immersion objective NA 1.4. A 488 nm laser was used for excitation and images were captured at a time interval of 5-16 s for 8-15 min. The longer time interval (16 s) was required when imaging the entire neuromast, including the hair cell bodies. Timelapses were acquired using the Zeiss Zen software.

### Image Analysis

Image analysis was done in ImageJ. Prior to quantification, the timelapses requiring bleach correction were processed using the ‘Bleach correction’ function in ImageJ by the simple ratio method. The timelapses were registered using the ‘Stackreg’ plugin in ImageJ by the rigid body method.

The fluorescence images shown in the figures were averaged over 4-10 frames using the ‘Walking Average’ function in ImageJ.

### Standard Deviation Projections

To obtain the standard deviation images of calcium activity, we cropped the cells and used the Z project function to create a standard deviation projection of the timelapse for the time period where activity was observed. The lookup table was changed to Rainbow RGB for better visualization and the contrast adjusted to be the same for all images in a panel to ensure a fair comparison.

### Difference Images

To obtain the difference images shown in Figure 6B, we used the ‘Stack Difference’ function in ImageJ using a 5 frame window. The resulting stack was overlayed on the original fluorescence image to create a composite. Each image shown is one frame of the difference stack where a transient was first seen.

### Temporal color-coded projections

The cells were cropped, and the time period of interest was extracted by duplicating the frames into a new stack. It is essential to have a short time period to get a clear temporal color code. Otherwise, the colors will overlap and merge in a longer timelapse. We used 25 s time periods and used the ‘Temporal-Color Code’ function in ImageJ to create the images and corresponding color bars in Figure 1C and 7B.

### Line Intensity Plots

The line intensity plots for Figure 3C were plotted using the ‘Plot Profile’ function in ImageJ. Each line plot corresponds to the first frame where the transient was seen. In Figure 3B, the first frames where transients were seen were plotted for 15 stereocilia.

To calculate the timescale of transients, the stereocilia and precursors were manually outlined and the mean pixel intensity calculated per frame using the ‘Plot Z axis Profile’ function in ImageJ. One example of a stereocilium and precursor is shown in Figure 4C.

### Quantification of stereocilia activity

To quantify spontaneous calcium activity, we counted the number of active stereocilia per hair cell visually.

### Electron Microscopy

For direct freezing and freeze substituted samples were processed as previously published. Briefly, the temporal bones were removed from adult rats and placed in Medium 199 (ThermoFisher) followed by microdissection. The isolated utricle vestibular sensory organ then transferred onto gelatin support that is mounted on an aluminum specimen holder. The utricles surface was rapidly brought against the surface of a liquid nitrogen-cooled sapphire block using a Life Cell CF-100 impact freezing machine. Samples were first transferred into liquid nitrogen then into the Leica Biosystems AFS for freeze substitution. Tissues were submerged in 1.5% uranyl acetate in absolute methanol at -90 °C for 2 days and infiltrated with HM20 Lowicryl resin (Electron Microscopy Sciences) over 2 days at -45 °C. The resin was polymerized with ultraviolet light for 3 days at temperatures between -45 °C to 0 °C. Ultrathin sections at 70 nm were generated using a Leica ultramicrotome and collected onto 300 mesh hexagonal copper grids (Electron Microscopy Sciences). Thin sections were imaged using a Zeiss/Leo922 electron microscope equipped with an energy filter operated at 200kV.

---

**Figure S1:**
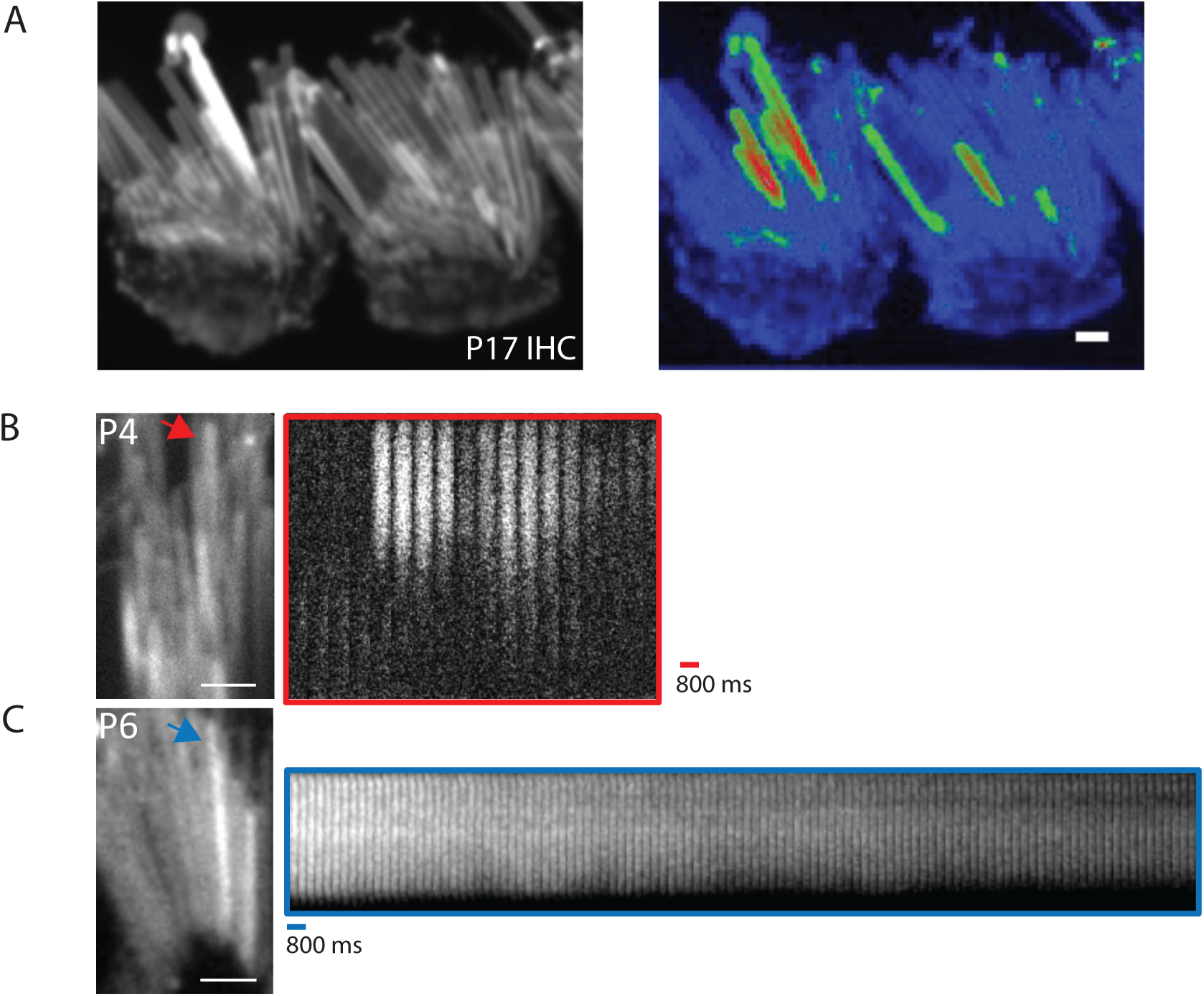
Individual calcium transients continue after the onset of hearing. **A)** Imaging membrane-localized GCaMP3 in acute tissue explants of the organ of Corti at P17 shows that these transients continue after the onset of hearing and MET. The left panel shows inner hair cells at P17 that were imaged at a time interval of 200 ms. The right panel shows the standard deviation projection of the timelapse, where calcium activity in individual stereocilia can be observed. **B)** A P4 utricle is shown with a kymograph of the stereocilium marked with a red arrow. The calcium transient peaks within a time frame of 800 ms. **C)** An example of a transient with a slow decay. Shown here is a P6 sacculus with a kymograph of the stereocilium marked with a blue arrow. Scale bars for all panels are 1 *μ*m.

